# Clock driven waves of Tbx6 expression prefigure somite boundaries

**DOI:** 10.1101/2023.11.09.566373

**Authors:** Olivier F. Venzin, Chloé Jollivet, Nicolas Chiaruttini, Olga Rosspopoff, Clément Helsens, Luis G. Morelli, Koichiro Uriu, Andrew C. Oates

**Affiliations:** Institute of Bioengineering, School of Life Sciences, Swiss Federal Institute of Technology EPFL, Lausanne, Switzerland; BioImaging and Optics Core Facility (BIOP), School of Life Sciences, Swiss Federal Institute of Technology EPFL, Lausanne, Switzerland; Global Health Institute, School of Life Sciences, Swiss Federal Institute of Technology EPFL, Lausanne, Switzerland; Instituto de Investigación en Biomedicina de Buenos Aires (IBioBA) – CONICET – Partner Institute of the Max Planck Society, Polo Científico Tecnológico, Argentina; Graduate School of Natural Science and Technology, Kanazawa University, Japan School of; Life Science and Technology, Tokyo Institute of Technology, Japan

## Abstract

The segmented body plan of vertebrates is established during embryogenesis by periodic and sequential formation of multi-cellular structures called somites. Somitogenesis is an example of patterning by a biological oscillator, the segmentation clock, which manifests as traveling waves of oscillating Hes/Her gene expression, reiterating during the formation of each^1–3^. How these waves are converted into the striped Mesp gene expression pattern that prefigures morphological somite boundaries^4–8^ remains unclear. Here, we image this conversion in real-time at single-cell resolution in zebrafish, using light-sheet microscopy of a novel reporter of Tbx6, a key activator of Mesp expression. We observe cellular oscillations and kinematic waves of Tbx6 expression that are driven by Hes/Her genes. Tbx6 waves arrest precisely in boundary cells that eventually express Mesp, thereby prefiguring the Mesp pattern, whereas Hes/Her waves do not. Although Hes/Her oscillations began before somitogenesis^9–11^, the first Tbx6 wave defines the boundary cells of the anterior-most somite, forming the head-trunk interface. Our findings imply that Tbx6 acts as a genetic clutch, converting Her/Hes pacemaker waves into Mesp stripes. We propose that this clock design shields the pacemaker from external perturbations, allowing flexible and robust patterning, making it of interest for organoids and tissue-engineering.

## Main

One of the major events of vertebrate embryonic development is the segmentation of the body axis into transient blocks of tissue, called somites, that ultimately give rise to vertebrae, ribs, and skeletal muscles of the adult body. Somites bud off from the presomitic mesoderm (PSM) by the sequential formation of new morphological boundaries. In the anterior PSM, sharp stripes of *mesp* gene expression (Mesp2 in mice, four *mesp* genes in zebrafish) are periodically expressed in the rostral part of presumptive somites, thereby establishing their rostro-caudal polarity^4,8^ and prefiguring the position of their future boundaries^7,12^. Mesp is one of a group of genes, such as *papc*^13,14^, expressed with a similar pattern that likely combine to effect the morphological transition of a somite boundary. New somite boundaries form between two rows of cells: the “anterior row” of a presumptive somite, consisting of cells located at the anterior edge of the stripes of *mesp* expression (Mesp-positive), and the “posterior row” of the previous somite which consists of Mesp-negative cells (Fig. 1b)^12,15^.

**Figure 1:**
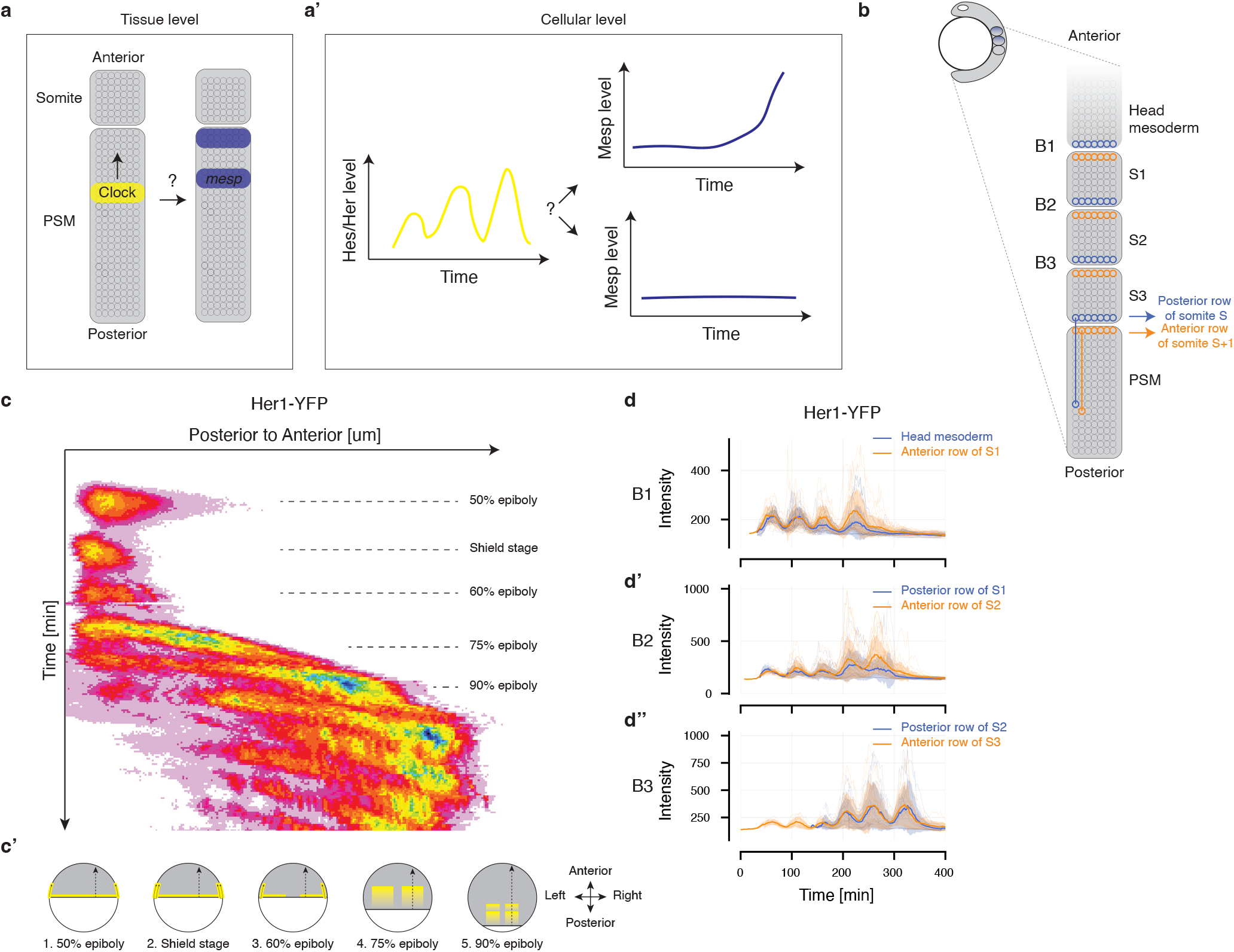
Waves of Her1-YFP do not arrest at a position prefiguring the *mesp* pattern. **a-a’,** How is the clock’s signal converted into spatially periodic pattern at tissue- (**a**) and cellular-level (**a’**)? **b,** Schematic of zebrafish embryo during early somitogenesis. Somite boundaries form between Mesp-positive cells in anterior row of presumptive somite (orange) and Mesp-negative cells in posterior row of previous somite (blue). Cells forming somite boundaries back-tracked through PSM (blue and orange dashed lines). **c,** Kymograph of the Her1-YFP signal during segmentation clock onset and formation of first somites (n= 3 embryos, 3 independent experiments). **c’** Schematic of location of first waves of Her1-YFP (yellow) during epiboly (Supplementary Discussion 1). Arrows indicate positions of the lines of interest used for kymograph in **c**. **d-d’’,** Mean and standard deviation of Her1-YFP intensities in thick line and shade, respectively. Signals from different embryos were temporally aligned using time of epiboly completion. Cells back-tracked from the head mesoderm, the posterior row of somites 1 and 2 (blue) and the anterior row of somites 1, 2 and 3 (orange), respectively (n = 29, 16, 15, 26, 30, 14 cells from 3 embryos, 3 independent experiments).

The T-box gene Tbx6, which is required for the expression of *mesp* and *papc*^12,14,16^ and for the formation of morphological somite boundaries^17,18^, shows a spatially periodic pattern of expression that defines the *mesp* expression pattern^6,19^. The periodic pattern of *mesp* expression is also thought to be regulated by the segmentation clock, which manifests as spatiotemporal waves of Hes/Her gene expression travelling anteriorly in the PSM and reiterating during the formation of each somite^1,3^. Indeed, when Hes/Her kinematic waves are abolished (e.g. in *her1;her7* zebrafish double mutants or in Hes7-null mouse embryos), the *mesp* pattern is disrupted and the resulting somite boundaries are defective^20–24^, suggesting that Hes/Her waves control which cells in the tissue will take the Mesp cell fate. Studies in mice and zebrafish have proposed plausible mechanisms to explain the conversion of the temporal signal from the clock into a spatially periodic pattern of gene expression, but the dynamics of molecular players was inferred from static snapshots of in mRNA situ hybridization and immunostaining^6,25^. How the temporal signal of the Hes/Her genes is converted into the spatial pattern of *mesp* expression, that ultimately prefigure the position of future somite boundaries, has never been observed in live embryos and therefore remains elusive, both at the tissue- (Fig. 1a) and at the cellular-level (Fig. 1a’). To address this question, we first followed cells in the PSM until they formed a given somite boundary, comparing the Hes/Her signal of Mesp-positive and Mesp-negative boundary cells, in an attempt to find a signature distinguishing the two cell populations.

### The onset of the segmentation clock

While comparing Hes/Her signals between Mesp-positive and Mesp-negative cells could be done for any somite boundary in the embryo, the first somite boundary is of particular interest for three reasons. First, this boundary marks the transition from head to trunk identity in the vertebrate embryo and has been an important subject of comparative zoology and evo-devo, as it plays a central role in defining the body plan of vertebrates^26^. Second, the onset of the zebrafish segmentation clock has only been studied using mRNA *in situ* hybridization^10,27^ and a dynamic understanding of this process is currently lacking. Third, several cycles of Hes/Her gene expression were reported to precede the onset of *mesp* expression and somite formation in mice, chick and zebrafish^9–11,27^. This suggests that several early waves of Hes/Her expression do not instruct cells to take a Mesp cell fate during epiboly. Therefore, imaging the onset of the segmentation clock allows the comparison, without any additional perturbation, of Hes/Her waves that are, or are not, involved in producing the Mesp pattern.

We injected embryos carrying Her1-YFP^3^ and Mesp-ba-mKate2^28^ transgenes at 1-cell stage with H2B-mCerulean mRNA to visualize nuclei, which served as markers for cell tracking and allowed us to unambiguously identify somite boundaries^11^ and the identities of cells that compose them (Fig. 1b). Embryos were imaged, using light-sheet microscopy, from the stage of 30% epiboly (4.7 hpf), before the onset of *her1* expression^27^, until mid-somitogenesis (Supplementary video 1). Kymographs generated from maximum-intensity projection of elliptically transformed timelapses (Supplementary video 2 and Methods) showed that three short-range waves of Her1-YFP expression traveled from the margin towards the animal pole (Fig. 1c), including cells fated to become neural ectoderm (Fig. 1c’). The fourth wave of Her1-YFP expression at 75% epiboly traveled further, restricted to the involuted mesodermal cells forming the PSM (Fig. 1c-c’). These results are the first description of Her1 dynamics during the onset of the zebrafish segmentation clock (Supplementary Discussion 1).

To compare the Her1-YFP signals of Mesp-positive (Fig. 1b, Anterior row of somite S+1) and Mesp-negative (Fig. 1b, Posterior row of somite) cells forming a given somite boundary, we selected cells forming the first three somite boundaries (Fig. 1b): The first, anterior-most somite boundary (B1) which separates the head mesoderm from the trunk mesoderm; the boundary between somite 1 and 2 (B2); and the boundary between somite 2 and 3 (B3). We then tracked these boundary cells back in time to obtain their Her1-YFP and their Mesp-ba-mKate2 signals. Cells forming the boundary B1 displayed on average 4 cycles of Her1-YFP expression as they were advected along the PSM (Fig. 1d). As each oscillation corresponds to a spatiotemporal wave of Her1-YFP expression, this suggests that the Mesp cell fate could be instructed by the wave number 4. Cells forming the boundary B2 or B3, displayed on average 5, respectively 6, oscillations of Her1-YFP expression (Fig. 1d’-d’’), implying that the fifth and sixth waves of Her1-YFP expression may instruct the Mesp cell fate of cells forming the second and third somite boundaries, respectively. We extrapolate that in zebrafish, the n^th^ somite boundary is prefigured by the wave number n+3. This provides the map between kinematic waves and the somite boundaries they prefigure in zebrafish.

Strikingly, cells located at the posterior of the head mesoderm, which are Mesp-negative, displayed on average the same number of Her1-YFP oscillations as boundary cells forming the anterior row of the first somite (Fig. 1d). This means that the waves of Her1-YFP expression did not stop at the first somite but continued to travel anteriorly in the head mesoderm. Similarly, cells forming boundaries B2 and B3 displayed respectively the same number of Her1-YFP oscillations, regardless of which side of the boundary they were located. These results suggest that the Mesp cell fate of boundary cells is not discriminated by the Her1-YFP pattern and confirms, with a cellular resolution, that the arrest of Her1 oscillations does not cause somite boundary formation^25^.

We then asked whether we could find a signature that distinguished between Mesp-negative and Mesp-positive cells forming a given somite boundary. Tbx6 is a good candidate, because it is required for Mesp expression^12,29^, and it spatially defines the Mesp pattern^6,19^. The existing Tbx6 transgenic line expresses a stabilized fluorophore preventing observation of short-term dynamics^30^. To overcome this limitation, we developed a new line where Tbx6 is fused with mNeonGreen (Extended Data Fig. 1), and we imaged the dynamics of the nuclear-localized Tbx6-mNG fusion protein at single-cell resolution throughout somitogenesis.

### Waves and oscillations of Tbx6-mNG

We imaged zebrafish embryos carrying Tbx6-mNG and Mesp-ba-mKate2 transgenes, injected with H2B-mCerulean mRNA at 1-cell stage, from the stage of 30% epiboly (Supplementary video 3). Kymographs generated from maximum-intensity projection of elliptically transformed timelapses (Supplementary video 4 and Methods) showed more than 13 waves of Tbx6-mNG expression traveling anteriorly in the PSM from 50% epiboly until mid-somitogenesis (Fig. 2a, arrowhead). To determine if these traveling waves of Tbx6-mNG expression are underlain by cellular oscillations, we back-tracked cells located within the first three somites of the embryo to record their dynamics as they travel along the PSM (Fig. 1b). Each individual cell displayed two to three oscillations of Tbx6-mNG (Fig. 2b-b’’). This result shows that coordinated cellular oscillations underlie tissue-level waves of Tbx6-mNG expression. To our knowledge, this is the first time that waves and oscillations of Tbx6 expression are reported in any species.

**Figure 2:**
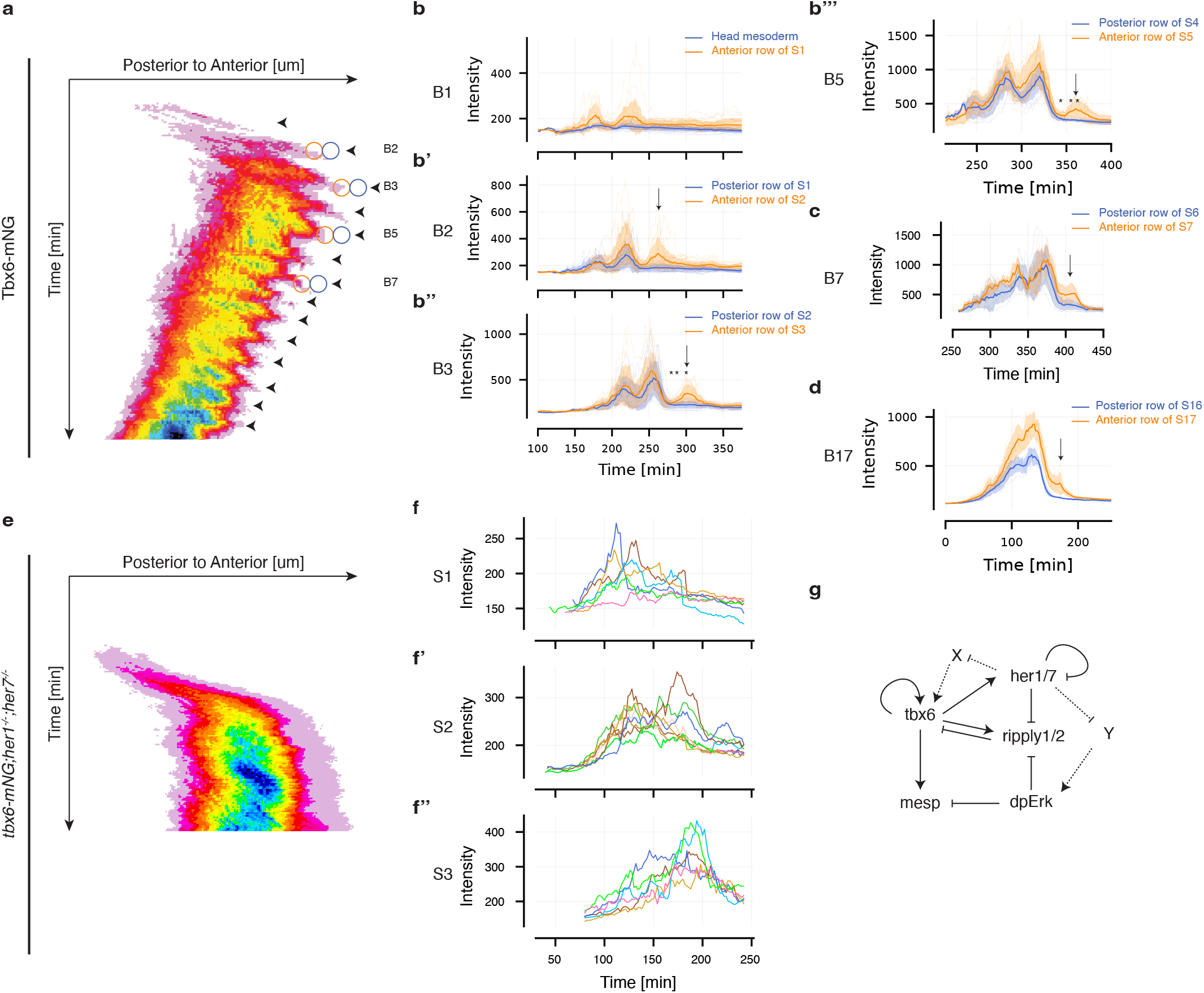
Clock-driven waves of Tbx6 arrest at position prefiguring the expression of *mesp*. **a,** Kymograph of Tbx6-mNG signal during early somitogenesis (n= 3 embryos, 3 independent experiments). Arrowheads indicate Tbx6-mNG travelling waves. Orange and blue circles mark positions of cells forming boundaries B2, B3, B5 and B7 at times shown by arrows in **b’-d**. **b-d,** Mean and standard deviation of Tbx6-mNG intensities displayed in thick line and shade, respectively. Signals from different embryos were temporally aligned using time of epiboly completion. **b-b’’’,** Cells back-tracked from the head mesoderm, located in posterior end of somites 1, 2, and 4 (blue) and in anterior end of somites 1, 2, 3 and 5 (orange), respectively (n = 26, 19, 14, 12, 26, 22, 17, 13 cells from 3 embryos, 3 independent experiments). **c-d,** Cells back-tracked from boundaries 7 and 17, respectively (n = 12 and n=10 cells from two embryos. Tbx6-mNG traces in cells forming boundaries 7 and 17 in other embryos shown in Extended Data Fig. 3). Arrows in **b’-d** indicate the extra cycle of Tbx6 that distinguishes Mesp-positive (orange) from Mesp-negative cells (blue). Stars in **b’’ and b’’’** mark timepoints represented in Extended Data Fig. 5 and Fig. 3, respectively. **e**, Kymograph of Tbx6-mNG signal in *her1;her7* double mutants (n= 3 embryos, 3 independent experiments). **f-f’’,** Individual Tbx6-mNG traces in cells located in somites 1, 2 and 3 of a *her1;her7* double mutant embryo (n = 6, 8, 6 cells respectively). Traces from another embryo shown in Extended Data Fig. 3c. **g,** Schematic of updated gene regulatory network controlling specification of Mesp cell fate.

### Her1/7 loop indirectly drives waves of Tbx6

Before investigating whether there is a signature in Tbx6 signals that can distinguish between Mesp-positive and Mesp-negative cells forming a somite boundary, we asked whether Tbx6 oscillations are driven by the known Her1/Her7 negative feedback loop of the segmentation clock, or whether they constitute an independent oscillator. To do so, we imaged Tbx6-mNG in *her1;her7* double mutants (Supplementary video 5). In zebrafish, *her1* and *her7* constitute the core of the clock and *her1;her7* double mutants are characterized by a disrupted *mesp* pattern and defective somite boundaries along the entire body axis^20,23^. Tbx6-mNG^+/−^;*her1^−/−^;her7^−/−^*embryos were injected with H2B-mCerulean mRNA at 1-cell stage for cell tracking. Waves of Tbx6-mNG expression were abolished in *her1;her7* double mutants (Fig. 2e). We back-tracked cells located within the region corresponding to the first three somites. Consistent with the absence of Tbx6-mNG waves, these cells showed a fluctuating Tbx6-mNG expression profile, but did not exhibit oscillations (Fig. 2f-f’’, Extended Data Fig. 3). Together, these results imply that the Her1/Her7 loop drives oscillations and kinematic waves of Tbx6-mNG expression. We then investigated whether the Her1/Her7 loop drives Tbx6 oscillations via a direct repression of *tbx6* expression. Although Her binding to gene regulatory sequences has been tested *in vitro*^31,32^, the binding of Her1 or Her7 to regulatory regions in the embryo remains unknown. To assess whether Her1 and Her7 directly bind to *tbx6*, we performed CUT&Tag^33^ using anti-GFP antibodies to address the genomic binding landscape of the fusion proteins Her1-YFP and Her7-YFP from their respective transgenic lines, as no antibodies against Her1 and Her7 currently exist. As a positive control, we confirmed that Her1 and Her7 are bound to intergenic regions in the proximity of known targets, such as their own chromosomal locus and that of *deltaC*^32,34^ (Extended Data Table 1 and 2, Extended Data Fig. 4a-b). However, no Her1 nor Her7 binding sites were found in the proximity of the *tbx6* gene (Extended Data Fig. 4e), suggesting that Her1 and Her7 do not drive Tbx6 oscillations by directly repressing *tbx6* expression.

Consistent with a recent report of the repression of *ripply1* and *ripply2* expression by Her1^25^, we found that Her1 and Her7 bound to the promoter regions of *ripply1* and *ripply2* (Extended Data Table 1 and 2, Extended Data Fig. 4c-d). As Ripply1/2 induce the degradation of Tbx6 proteins^6,7,19,35^, there is a possibility that the Her1/Her7 loop drives Tbx6 oscillations indirectly by regulating *ripply1/2* expression. However, Tbx6 oscillations are observed more posteriorly in the PSM than the expression of either *ripply*1 or *ripply2*^19,25,36^. In addition, Ripply1/2 proteins induce a long-lasting repression of Tbx6^25^. Combined, these facts argue that neither the transcriptional nor the post-transcriptional regulation of Tbx6 by Ripply1/2 proteins can explain the oscillations of Tbx6 in the posterior PSM. Thus, we conclude that Tbx6 oscillations are driven indirectly by the Her1/Her7 loop via a novel, yet-to-be-identified regulatory player.

### Tbx6 traces distinguishes boundary cells

To investigate the role of clock-driven waves and oscillations of Tbx6 in Mesp cell fate specification, we compared Tbx6 traces between cells that eventually become Mesp-positive or Mesp-negative by back-tracking cells forming the first three somite boundaries (Extended Data Fig. 2a-b). Cells located within the head mesoderm, which do not express Mesp (Extended Data Fig. 2c, d-d’’), did not express Tbx6 (Fig. 2b, Head mesoderm). This cell population displayed four Her1-YFP oscillations (Fig. 1d), indicating that the first oscillations of Her1-YFP are independent of Tbx6. Strikingly, cells forming the anterior boundary of somite 1, which are located just posteriorly to the cells in the head mesoderm, express Mesp (Extended Data Fig. 2c, d-d’’) and displayed two cycles of Tbx6-mNG expression (Fig. 2b, Anterior row of S1). This indicates that the onset of Tbx6 oscillations and the corresponding formation of the Tbx6 domain spatially defines the first *mesp* stripe, which itself prefigures the position of the boundary separating the head and the trunk mesoderm.

Mesp-positive cells forming the anterior row of somite 2 displayed one more oscillation of Tbx6-mNG than Mesp-negative cells forming the posterior row of somite 1 (Fig. 2b’ and Extended Data Fig. 2c’, e-e’’). Similarly, Mesp-positive cells forming the anterior row of somite 3 displayed one more oscillation of Tbx6-mNG than Mesp-negative cells forming the posterior row of somite 2 (Fig. 2b’’ and Extended Data Fig. 2c’’, f-f’’). To extend these observations, we back-tracked cells forming the fifth, the seventh and the seventeenth somite boundaries. We observed that, although the peak-to-trough difference of the Tbx6-mNG oscillatory traces decreased over developmental time, the extra cycle of Tbx6-mNG continued to reliably discriminate cells in the Mesp-positive anterior row from the Mesp-negative posterior row (Fig. 2c-d and Extended Data Fig. 3a-b). These results, together with the observation of more than 13 waves of Tbx6-mNG expression, suggest that the number of oscillations of Tbx6 expression distinguishing between Mesp-positive and Mesp-negative cells forming a somite boundary is a general feature of somitogenesis. Given that Tbx6 is a required activator of Mesp expression^12,16,29,37^, our results strongly suggest that the extra oscillation of Tbx6 expression instructs cells to a Mesp-positive versus a Mesp-negative cell fate. Since each cellular oscillation corresponds to a spatiotemporal wave of Tbx6-mNG expression at the tissue-level, Tbx6 waves should arrest at a location that precisely prefigures *mesp* expression. We confirmed this by visualizing the positions of individual Mesp-negative and Mesp-positive cells in Tbx6-mNG kymographs (Fig. 2a, blue and orange circles, respectively).

### The periodic pattern of Tbx6 expression

We next investigated whether these clock-driven waves of Tbx6-mNG expression are consistent with the periodic pattern of endogenous Tbx6 protein^19,25^? In zebrafish, immunostainings against Tbx6 show a periodic pattern characterized by a core domain of Tbx6 protein expression and the presence or absence of an anterior stripe of Tbx6 protein expression^19^. We term these two phases “Anterior stripe” and “Absence of anterior stripe”, respectively. The expression of Tbx6-mNG in our timelapses also showed the periodic appearance (Fig. 3a-a’, e-e’, white arrowheads) and disappearance (Fig. 3c-c’) of an anterior stripe of Tbx6 expression, directly comparable to the patterns revealed by anti-Tbx6 immunohistochemistry (Fig. 3b, d, f). Intensity profiles of Tbx6 along the AP axis of the PSM captured the presence (Fig. 3a’, b’, e’, f’, black arrowheads) and absence (Fig. 3c’, d’) of the anterior stripe of Tbx6 expression in both the Tbx6-mNG timelapses and the immunostaining against Tbx6, confirming that the Tbx6-mNG transgenic recapitulates the spatiotemporal expression of endogenous Tbx6 proteins.

**Figure 3:**
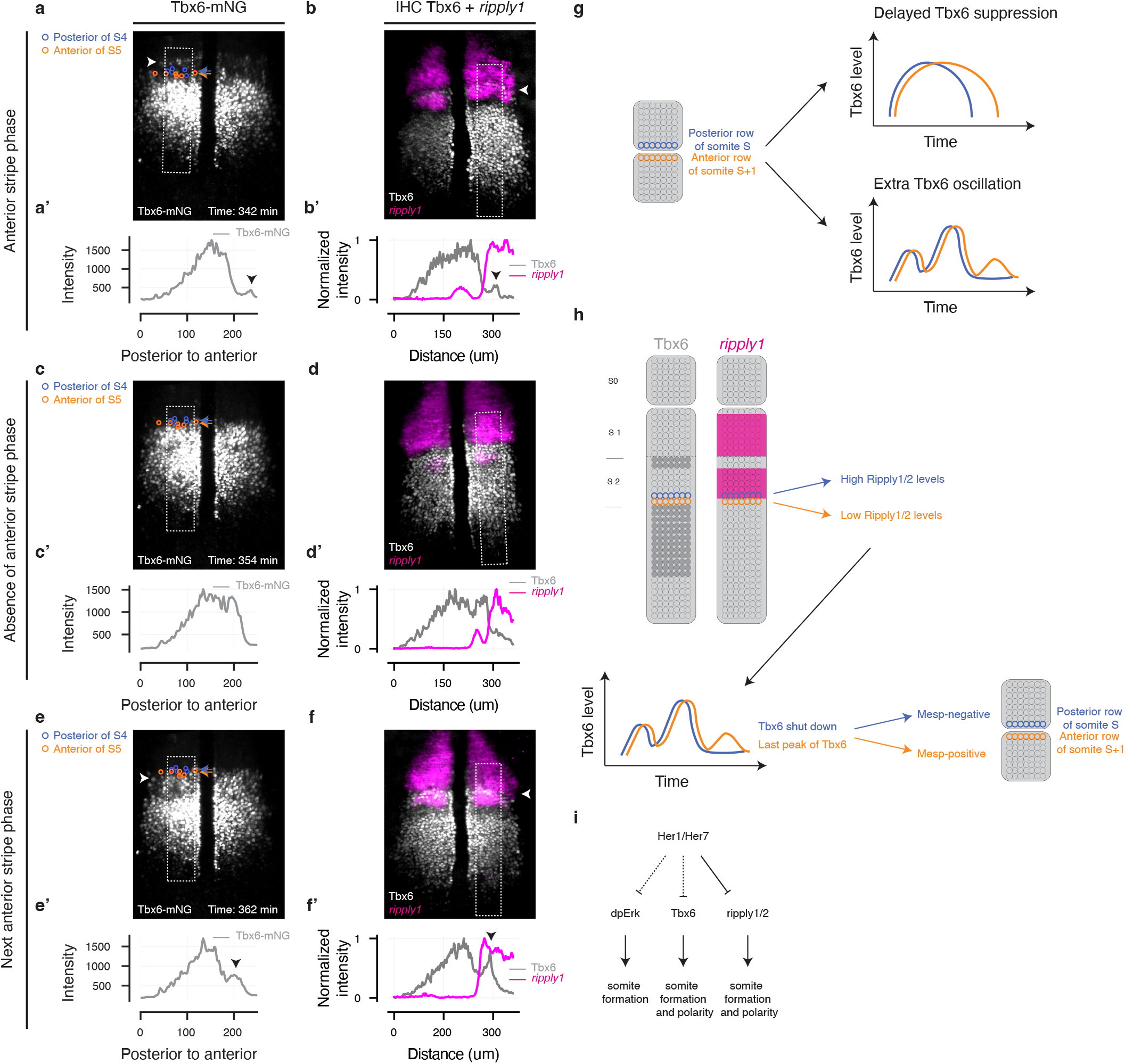
The anterior stripe of Tbx6 corresponds to the last cycle of Tbx6. **a, c, e,** Snapshots from Tbx6-mNG timelapse: the “anterior stripe” phase (**a)**, the “absence of anterior stripe” (**c)** and the next “anterior stripe” (**e**) phases, respectively. Blue and orange circles mark positions of cells located in the posterior row of somite 4 and in the anterior row of somite 5, respectively. **b, d, f,** Tbx6 immunostaining (grey) combined with *ripply1* mRNA *in situ* hybridization (magenta) of representative 4-somite stage embryos (n = 14 embryos). White boxes show areas of interest used to plot mean Tbx6 profiles along PSM (**a’-f’**). White arrowheads indicate anterior stripe of Tbx6-mNG expression (a, c), black arrowheads indicate signature of this stripe on intensity profiles (a’, c’). Tbx6-mNG patterns and profiles match Tbx6 immunostaining for all phases. **g**, Schematic of alternate mechanisms explaining formation of anterior stripe of Tbx6 expression: delayed suppression versus extra oscillation of Tbx6. **h,** Schematic of the proposed mechanism by which Ripply1/2 shut down oscillations of Tbx6 and precisely arrest waves of Tbx6 expression, leading to a sharp transition in the Mesp cell fate. **i,** The Her1/Her7 loop acts on multiple downstream effectors either directly (plain line) or indirectly (dashed line).

How are the discrete borders of Tbx6 proteins created in the anterior PSM? Previous models, based on fixed snapshots, proposed that the anterior stripe of Tbx6 expression was generated by Ripply1/2-induced degradation of pre-existing Tbx6 protein in a two-step process^19^: The first step is the degradation of Tbx6 protein in the anterior part of the core domain, except for its anterior-most part, resulting in the appearance of a trough separating the anterior stripe of persisting Tbx6 protein from its core protein domain. The degradation then propagates anteriorly, induced by increasing Ripply1/2 protein expression, which ultimately results in the disappearance of the Tbx6 anterior stripe. Since the anterior stripe of Tbx6 is thought to spatially control the expression of *mesp*^6,19^, this interpretation of the mechanism underlying the Tbx6 pattern implies that Tbx6 is first degraded in Mesp-negative cells forming the posterior row of one somite and later in the Mesp-positive cells forming the anterior row of the next somite (Fig. 3g, Delayed Tbx6 suppression). However, our finding that the expression of Tbx6 protein oscillates raises the possibility that the anterior stripe of Tbx6 corresponds to an extra oscillation (i.e. *de novo* expression) and not only from persisting Tbx6 proteins^19^.

To test this hypothesis, we followed, in a Tbx6-mNG timelapse, the positions of Mesp-positive and -negative cells eventually forming the boundary B5 at three time points corresponding to different phases of the Tbx6 pattern (Fig. 3a, c and e, blue and orange circles, respectively) and we compared their Tbx6-mNG expression at these phases (Fig. 2b’’’, stars). During the phase “Anterior stripe” that prefigured the Mesp pattern of the previous boundary (B4), cells that will eventually form the boundary B5 were located in the trough between the core domain and the anterior stripe of Tbx6-mNG expression (Fig. 3a, blue and orange circles). Consistent with their positions, the intensity of their Tbx6-mNG expression at that stage is low (Fig. 2b’’’, left star).

The anterior stripe prefiguring the boundary B4 then disappears, leaving only the core domain of Tbx6 expression which characterizes the phase “Absence of anterior stripe”. During this phase, cells that will eventually become Mesp-positive and form the anterior row of somite 5 were located at the edge of the Tbx6 core domain (Fig. 3c, orange circles) and displayed their last cycle of Tbx6-mNG expression (Fig. 2b’’’, middle star). During the next “Anterior stripe” phase, these cells were part of the anterior stripe of Tbx6 expression (Fig. 3e, orange circles) and were still in their last cycle of Tbx6-mNG expression (Fig. 2b’’’, right star). This implies that the duration of the last cycle spans these two spatial phases.

In contrast, cells located just anteriorly that will eventually be Mesp-negative and form the posterior row of somite 4 (Fig. 3c and e, blue circles), did not express Tbx6-mNG at these times (Fig. 2b’’’, middle and right stars). The same relationships were found by comparing the oscillatory traces and the positions of cells forming the boundary B3 (Extended Data Fig. 5 and Fig. 2b’’, stars). These results indicate that the anterior border of the Tbx6 broad domain and the anterior stripe of Tbx6, which are thought to spatially control the expression of *mesp*^6,19^, correspond to the extra oscillations of Tbx6 traces displayed by Mesp-positive cells forming the anterior row of somites (Fig. 3 and Fig. 2b’’-b’’’). Importantly, this also implies that the anterior stripe of Tbx6 expression arises with a contribution from *de novo* expression of Tbx6 (Fig. 2b’’-b’’’).

### Ripply1/2 prevents the extra cycle of Tbx6

How do we incorporate this new observation in the current understanding of how Ripply1/2 proteins fix the anterior border of the Tbx6 protein domain^19,25^? Using combined whole-mount *in situ* hybridization for *ripply1* and immunostaining against Tbx6, we compared the spatial distribution of Tbx6 protein and *ripply1* expression (Fig. 3b-b’, d-d’, f-f’). During the first “anterior stripe” phase, which happens before the Mesp-positive cells forming the anterior row of somite 5 display their last cycle of Tbx6 expression (Fig. 2b’’’, left star), *ripply1* mRNA (Fig. 3b) and Ripply1/2 proteins^25^ were expressed in the trough between the core domain and the anterior stripe of Tbx6 protein expression, but their expression did not extend all the way to the Tbx6 core domain. Cells that will eventually display an extra cycle of Tbx6 and become Mesp-positive were located just anterior to the core domain of Tbx6 (Fig. 3a, orange circles), in a region devoid of Ripply1/2 proteins^25^ and *ripply1* mRNA (Fig. 3b), while cells that will not display an extra cycle of Tbx6 and become Mesp-negative were located slightly more anterior (Fig. 3a, blue circles) in a region where Ripply1/2 proteins^25^ and *ripply1* mRNA are likely expressed (Fig. 3b). The spatial relationship between the positions of cells forming somite boundaries (Fig. 3, Extended Data Fig. 5) and Ripply1 protein and mRNA expression domains means that future Mesp-positive and Mesp-negative cells are likely to experience different levels of Ripply1 protein. As Tbx6 expression has recently been shown to respond to different levels of Ripply1/2 proteins in a switch-like manner^25^, these results suggest that higher levels of Ripply1/2 proteins prevent cells from displaying an extra cycle of Tbx6, whereas cells located just posterior to the Ripply1/2 protein domain would show a last peak cycle of Tbx6 expression, which in turn induces Mesp expression^16,37^. The spatial pre-pattern of *mesp*^12,37^ is thus defined, via the location of the posterior-most expression of Ripply1/2 proteins, by the arrest of the wave of Tbx6.

## Discussion

The transition from head to trunk identity in the vertebrate embryo has been the subject of long-standing debate in embryology and evolution^38^. The first oscillations of the segmentation clock observed in mouse, chick and zebrafish before somite formation have appeared paradoxical, as they occur in tissue primordia that will not form somites^9–11^. There is no evidence connecting the oscillations to putative head metamerism^39,40^, and the role of these warm-up oscillations, if any, remains unclear. Our findings reveal that the transition from unsegmented head mesoderm to the patterning of segmented trunk mesoderm is marked by the onset of oscillatory Tbx6 expression, rather than a change in the segmentation clock (Fig. 1 and Fig 2). Tbx6 expression thus acts as a clutch to couple the already-running segmentation clock to the suite of genes necessary to cause the formation of somite boundaries. Control of Tbx6 onset may be related to early RA signaling^41^, but is beyond the scope of the current study.

Patterning tissues with oscillations is emerging as a common mechanism in development and regeneration^42^. In this paper, we describe how the Her1/Her7 loop of the segmentation clock acts as a pacemaker, spatiotemporally controlling the expression of the key effector Tbx6, that directly controls somite segmentation and the rostro-caudal polarity of somites (Fig. 2). We speculate that separating the pacemaker and effector genes allows the downstream effectors to be spatiotemporally modulated by external factors while shielding the pacemaker from these perturbations.

Our results suggest that the posterior-most protein expression of Ripply1/2 defines the position where the waves of Tbx6 arrest (Fig. 3). Interestingly, the expression of *ripply1/2* is regulated by dpErk^19,25^, the Her1/7 loop^25^ (Extended Data Table 1 and 2, Extended Data Fig. 4c-d) and Tbx6^36^. In a manner that remains elusive, the Her1/7 loop drives oscillations of both dpErk^43^ and Tbx6 (Fig. 2). Therefore, the Her1/7 loop spatiotemporally controls the expression of *ripply1/2* both directly, and indirectly via dpErk and Tbx6 (Fig. 3i). Periodic inhibition of Erk activity was recently shown to be sufficient to produce sequential segmentation^43^. However, this periodic inhibition of dpErk alone did not produce a striped Mesp pattern, and also failed to produce rostro-caudally polarized segments^43^. Our results suggest that the endogenous Mesp pattern is established by waves of Tbx6 expression as part of the complete somite structure.

How does the last peak of Tbx6 cause *mesp* to be expressed in a cell, when the total amount of Tbx6 expressed by a cell with or without the last cycle may be very similar (Fig. 2, 3), and the previous two oscillations did not activate *mesp*? One likelihood is that the timing of the last peak is the key, rather than the amount of Tbx6, which implies that some other factor is creating a window of opportunity for Tbx6 action on the *mesp* promoter. A candidate for this factor is dpErk activity, which is thought to be globally repressive for *mesp* expression^12,19^. The posterior-ward movement of the anterior limit of the dpErk gradient that spans the PSM may create a short temporal window of opportunity in which a short pulse of Tbx6 acts as the clock-driven trigger for creating the *mesp*-positive state^43,44^.

It was recently proposed that the clock converts its temporal signal into a spatial pattern by direct periodic inhibition of *ripply1/2* expression^25^. Our results reveal two additional mechanisms: Firstly, since Tbx6 directly activates the expression of *mesp*^12,16,37^, clock-driven oscillations of Tbx6 periodically activate the expression of Mesp. Secondly, as mentioned above, given that Tbx6 also activates *ripply1/2* expression^35,36^, clock-driven oscillations of Tbx6 periodically activate the expression of *ripply1/2.* The determination of the Mesp cell fate is thus controlled by a densely connected gene regulatory network (Fig. 2h). These results indicate that the Her1/Her7 loop acts through multiple downstream effectors to translate its temporal signal into a spatial pattern. Our cell-tracking experiments revealed that the amplitudes of Her1-YFP oscillations were highly variable, with a coefficient of variation of 0.24 for the last two peaks. For target genes in the network that are sensitive to Her protein levels, this may introduce significant fluctuations, thereby decreasing the precision of the received temporal signal. We speculate that having multiple channels of control ensures accuracy in the Mesp patterning, which is crucial for the integrity of somite boundaries. In mice, oscillations in Notch activity are thought to periodically activate Mesp2 expression^6,45^. It remains unclear whether this is the only mechanism by which the mouse clock translates its temporal signal into a spatially periodic pattern, or whether there are multiple channels at work, as is the case in zebrafish.

In conclusion, our work reveals two design principles of the segmentation clock: the use of multiple channels of control and a separation between the pacemaker and its downstream effectors. These two properties might provide robustness and flexibility in a patterning system. Aside from providing insights into patterning in naturally occurring developing systems, these design principles may find applications in synthetic morphogenesis such tissue engineering and organoids.

## Methods

### Animals

Wildtype (AB and TL), mutants (*her1^−/−^;her7^−/−^*)^23^ and transgenic zebrafish Tg(*her1:her1-yfp*)^3^, Tg(*mesp-ba:mesp-ba-mkate2*)^28^, Tg(*tbx6:tbx6-mneongreen*)(this study; Extended Data Fig. 1) were maintained according to standard procedures on a 14h-10h light cycle at the EPFL fish facility which has been accredited by the the Service de la Consommation et des Affaires Vétérinaires of the canton of Vaud – Switzerland (VD-H23). Embryos were produced by natural spawning and staged according to (Kimmel *et al.* 1995)^46^. Heterozygous transgenic embryos were used for experiments. Embryos were incubated at 28.5°C in E3 medium (5mM NaCl, 0.17mM KCl, 0.33mM CaCl2, 0.33mM MgSO4, pH adjusted to 7.3 with Tris-HCl pH 7.5 1M). Embryos imaged during mid-somitogenesis were shifted to a 19.5°C incubator after shield stage.

### Generation of Tg(*tbx6:tbx6-mneongreen*) line

The transgenic line *TgBAC(Tbx6:Tbx6-mNeongreen)* was generated using BAC recombineering as previously described^3^. The BAC DKEY-232O23 containing Tbx6 locus as well as proximal genes sequences was used as starting material to create a functional fusion protein between the coding regions of Tbx6 and mNeonGreen. We inserted a 10 amino acid linker sequence (GGSGGSTASG) replacing the Tbx6 STOP codon, using the following recombineering primers:

Tbx6-HAL: ATGCTCATGGTCAGGGCTCTTACTTTGACCTTGGAGGACGAACAGTGTTCGGAGGAAGCGGAGGAAGC, and

Tbx6-HAR: GACATGTCACTTGCCAGCTCTTGACTTGCTTGTCATGTTTCTGTTTCTCTCGTCAGTCAGTACCGTTCG.

A total region of 23 847 bp containing TBX6 locus (8532 bp upstream *tbx6* 5’UTR and 3047 bp downstream the 3’UTR) fused with mNeonGreen was then subcloned into a pShave vector containing I-SCEI Meganuclease cutting sites using the following subcloning primers: Subcl-HAL: CTCAGACAGAATGAACGCTATAAAAGCAGTGCCTGATAAGACATGTAGAGCTATAGTGTCACCTAAATC, and

Subcl-HAR: (GCTGAGAAAAGACACTCCACATGATGGAACATTGGTTGAATATATTTTATCCCTATAGTGAGTCGTATTA).

40 pg of the final construct was then co-injected together with the I-SceI meganuclease enzyme (NEB) into one-cell stage AB wild type embryos. Founders were identified by outcrossing putative transgenic adult fish to WT animals and screening for expression of green fluorescence into the F1 embryo generation at 10 somites stage. The transgenic line was generated from a single founder transmitting as a single Mendelian locus and stably expressing mNeongreen after 3 generations.

*In situ* hybridization for *mNeonGreen* and *tbx6* riboprobes on 10 ss embryos revealed that the pattern of *tbx6-mNeonGreen* expression in *Tg(tbx6:tbx6-mNeongreen)*-positive embryos was similar to the endogenous expression of *tbx6* in their negative siblings (Extended Data Fig. 1). In addition, the expression of Tbx6-mNG in light-sheet timelapses showed the distinct patterns of Tbx6 expression (Fig. 3a, c, e)^19,25^ confirming that the Tbx6-mNG transgenic recapitulates the spatiotemporal expression of endogenous Tbx6 proteins. We assessed whether the Tbx6-mNG transgene rescues the *tbx6*^−/−^ phenotype by outcrossing *tbx6^−/−^*and *Tg(tbx6:tbx6-mNeongreen);tbx6^+/−^* adults. Using *in situ* hybridization against *xirp2a* to visualize the myotome boundaries, we found that somites formed in 100% (27/27) of *Tg(tbx6:tbx6-mNeongreen)*-positive embryos while somites only formed in 50% (15/30) of their negative siblings, confirming that the Tbx6-mNG transgene rescues the *tbx6^−/−^* mutants and is therefore functional.

We counted the same number of myotome boundaries at the proctodeum in *Tg(tbx6:tbx6-mNeongreen)*-positive embryos and in their negative siblings (17 boundaries), suggesting that the period of somitogenesis is not affected by the transgene. *In situ* hybridization staining against *xirp2a* revealed minor sporadic defects in the myotome boundaries in about half the embryos (14/26). These embryos make, on average, less than five defects (4.57) along their entire axis but no defects were observed in the first seven somite boundaries. Embryos imaged during mid-somitogenesis were stained for *xirp2a* to ensure that they had no myotome defects.

### Whole mount chromogenic *in situ* hybridization

Chromogenic *in situ* hybridization staining for *xirp2a*^47^ was performed according as previously described^48^ to assess the integrity of myotome boundaries. Myotome boundaries of embryos were scored on a binocular microscope (Olympus SZ61) and documented with a RGB camera (Olympus DP22). We confirmed the genotype of Tbx6-mNG;her1^−/−^;her7^−/−^ embryos after light-sheet timelapses by assessing the phenotype of their myotome boundaries.

### Whole-mount combined immunostaining and *in situ* hybridization

For simultaneous detection of *ripply1* mRNA and Tbx6 protein, we combined *in situ* hybridization and immunostaining. Digoxigenin-label RNA probes for *ripply1* were synthesized as previously described^48^. We first performed fixation, dehydration, hybridization and post-hybridization washes on 4 somite stage embryos as previously described^48^. Prior to hybridization, embryos were permeabilized with a 15 min incubation in Mili-Q water. Embryos were incubated at 4°C overnight with the primary antibody mouse IgG1 Anti-Tbx6 (home-made, clone A83, dilution 1:500) in 2% Blocking Reagent (Roche) and 5% sheep serum (Sigma) diluted in maleic acid buffer with 0.1% tween-20 (MABT). Embryos were washed 6 times with MABT and were then incubated with anti-DIG AP-conjugated Fab fragment (Roche, 11093274910), Alexa-Fluor 488 goat anti-Mouse IgG1 antibody (ThermoFisher A-21121, Dilution 1:1000) and DAPI (ThermoFisher, 1:1000 dilution) in 2% Blocking Reagent (Roche) and 5% sheep serum (Sigma) diluted in MABT. After 6 washes in MABT, *ripply1* mRNAs was detected using FastRed (Sigma). Whole-mount embryos were imaged using the Viventis LS1 light-sheet microscope with 3um z-slices and 3.3 um laser beams. Stack images were converted to xml/HDF5 using the Fiji plugin BigDataViewer^49^, elliptically transformed using BDV-playground (https://github.com/bigdataviewer/bigdataviewer-playground) and maximum-intensity projected. LOIs were used to measure the profile of Tbx6 or *ripply1* signals along the PSM. For comparison between different signals, the latter were normalized using custom Python scripts.

### CUT & TAG

We performed Cut&Tag as previsouly described^33^ on *Tg(her1:her1-YFP)* or *Tg(her1:nls-mkate2,her7:her7-YFP*) embryos. Briefly, 120 bud stage embryos per replicate were dechorionated using 1mg/ml Pronase (Roche) and dissociated into single cells with wash buffer (20 mM HEPES, pH 7.5, 150 mM NaCl, 0.5 mM spermidine and a protease inhibitor (Roche; one complete tablet per 50 ml added fresh)) by pipetting up and down using a 200 µl pipette. Low binding 2mL tubes were used to limit cells sticking to the wall. The dissociated single cells were then washed three times and bound to activated Concanavalin A-coated magnetic beads. Cells were resuspended into 100 µl of antibody buffer (wash buffer containing 0.05% (wt/vol) digitonin (Dig), 2 mM EDTA and 0.1 % BSA). Antibody at 1mg.ml was added to a final concentration of 1:50 and cells were incubated overnight at 4°C on a nutator. The next day, cells were resuspended in 100 µl of Dig wash buffer (wash buffer containing 0.05% (wt/vol) Dig) containing the secondary antibody at a final concentration of 1:100 and incubated for 60min at room temperature. After incubation, beads/cells were washed three times with Dig-wash buffer to remove unbound antibody, followed by resuspension and incubation of beads/cells in 100 µl Dig-300 wash buffer (Dig wash buffer at 300 mM NaCl) with pA-Tn5 adapter complex (1:200 dilution), at room temperature for 1 h on a nutator. After two washes with Dig-300 wash buffer, beads/cells were resuspended in 100 µl tagmentation buffer (Dig-300 buffer containing 10mM Mgcl2) and incubated at 37°C for 1 hour. Tagmentation reaction was stopped by adding exactly 3.3µl 0.5M EDTA, 1µl 10% SDS and 1µl 20mg.ml Proteinase K followed by 60 min incubation at 55°C shaking at full speed (1300 rpm) to solubilize DNA fragments. Beads/cells were placed on a magnetic stand and the supernatant containing DNA fragments was transferred into a new tube for DNA purification (MinElute Qiagen, 28004). The CUT and TAG experiment was done for two biological replicates using antibodies anti-GFP (Chromotek, PABG1-100, 1:50), anti-YFP (Merck Millipore, MABE1906, 1:25 dilution), H2A.Z (Active Motif, 39013, 1:50 dilution) was used as a positive control to confirm a successful experiment and a secondary antibody anti-Mouse IgG (Abcam, ab46540, 1:50 dilution) was used as negative control to remove background in the following sequencing analysis.

Cut&Tag libraries for next generation sequencing were prepared by PCR reaction using the NEBnext HIFI 2x PCR master mix and indexed i5 and i7 10µM primers as previously described^50^. Samples were then cleaned up using the DNA Clean and concentrator-5 kit from Zymo. Libraries were pooled together and paired-end 76bp sequencing was done using the Illumina NextSeq 500 instrument for Tg(her1:her1-YFP) samples and the MiSeq instrument for Tg(her1:nls-mkate2,Her7:her7-YFP) samples.

Paired reads were quality checked by FastQC (version 0.11.9) and aligned to the zebrafish genome assembly (GRCz.11 toplevel, Ensembl release 109) using Bowtie2 (version 2.5.1)^51^. The resulting sam files were then converted into bam files, sorted and indexed using samtools (version 1.17) and visualized with the Integrated Genomics Viewer (IGV version 2.16.0). Overlapping peak between the biological replicates was done using bedtools intersect (version 2.31.0) and peak calling was performed using macs2 callpeak (version 2.2.7)^52^ with the following parameters: -g 1.5e+9 -f BAMPE -q 0.01 --keep-dup all. Target genes near the peaks were manually annotated using IGV based on the same zebrafish genome assembly used for mapping.

### Light-sheet imaging

For light-sheet imaging, we used a Viventis LS1 microscope (Viventis Microscopy Sárl, Switzerland) equipped with a Hamamatsu Orca-Fusion BT CMOS camera, a CFI75 Apochromat 25X, NA 1.1 detection objective (Nikon), 405nm, 488nm, 515nm and 568nm laser lines and scanned gaussian beam light sheets with thickness of 3.3μm, respectively 2.2um, for timelapses that started during epiboly, respectively mid-somitogenesis. Tg(*her1-Venus;mesp-ba-mkate2*), Tg(*tbx6-mneongreen;mesp-ba-mKate2*) and Tg(*tbx6-mneongreen; her1^−/−^;her7^−/−^*) embryos were injected at 1-cell stage with 0.1 ng of H2B-mCerulean mRNA, *in vitro* transcribed from a pCS-H2B-Cerulean plasmid (Addgene, 53748), for nuclei visualization and tracking. Embryos were dechorionated and placed in an imaging chamber filled containing 2% low-melting agarose (Sigma) that was patterned to support, but not embed, embryos. The chamber was filled with 1ml of E3 medium and was maintained at 28.5 °C during acquisition. For imaging starting during epiboly, the agarose was patterned with spherical depressions with a trough pattern underneath so that the tail of the embryos can freely elongate. Embryos were mounted in the spherical depressions with the animal pole facing upwards. Every 2min, z stacks were acquired at 3um intervals. For imaging during mid-somitogenesis, embryos were mounted laterally with the yolk sitting in conical depressions patterned in the agarose and the medium was supplemented with 0.014% Tricaine to prevent muscle twitching during acquisition. Every 2min, z stacks were acquired at 2um intervals. Raw images were compressed using the compression system Jetraw (Dotphoton) and decompressed for image analysis.

### Image analysis

Every step of image analysis was performed using Fiji^53^. The basic processing steps (pre-processing, registration and cell tracking) of lightsheet timelapses were performed as previously described^54^. Briefly, timelapses were converted to XML/HDF5 files using BigDataViewer^49^ and registered using BigStitcher^55^. Boundary cells are identified using their H2BmCerulean and Mesp-ba-mKate2 signals and back-tracked in the PSM using their H2B-mCerulean signal either manually or semi-automatically with Mastodon v1.0.0-beta-20 (https://github.com/mastodon-sc/mastodon). Tracks obtained semi-automatically were manually curated to ensure that cells were accurately tracked. We used the Mastodon feature « Spot center intensity » to obtain the signals of interest from cell tracks.

As embryos are spherical during epiboly, timelapses were elliptically transformed using the BDV-playground (https://github.com/BIOP/bigdataviewer-biop-tools) to represent the curved surface of the embryo on a plane, as in a map projection. Briefly, an ellipsoid was fitted to the embryo and the dataset was transformed to be represented in spherical coordinates. As for a Mercator projection, the poles are distorted but the equator is preserved. Therefore, the equator of the ellipsoid was set as an anteroposterior line located at an equidistance between the two sides of the PSM. During epiboly, this line is perpendicular to the margin, passing through the shield. During somitogenesis, the line aligns with the notochord. We then projected the transformed dataset using a maximum-intensity projection (Supplementary Video 2 and 4).

Kymographs were obtained using the Fiji plugin “LOI interpolator” that interpolates lines of interest on maximum-intensity projections^3^. To obtain kymographs of embryos during epiboly, elliptically transformed timelapses were maximum-intensity projected and LOIs were traced from the margin of the blastoderm to the animal pole.

To visualize the positions of cells tracked with Mastodon on maximum-intensity projections of elliptically transformed timelapses, the same elliptic transformation was applied to the positions of cell tracks. The positions of cell tracks could then be displayed on the maximum-intensity projection of timelapses with the plugin TrackMate^56^.

### Data analysis

Cell traces were analyzed and plotted using custom script written in Python. Cell tracks from different embryos were temporally aligned using the time of epiboly completion as a reference.

### Statistical analysis

Statistical significance was assessed with independent two-sample t-test (two-tailed) using scipy.stats in Python. p-values lower than 0.05 were considered to indicate statistical significance.

## Acknowledgments

We thank current and past members of the Oates lab for scientific discussions and feedback on the manuscript. We thank Guillaume Valentin for his inputs and feedback over the entire course of the project.

## Author contributions

AO and OV conceived the project. AO supervised the project. CJ generated the transgenic line Tbx6-mNeonGreen. OV, CJ and OR performed experiments. OV and CJ tracked cells. OV and NC performed image analysis. OV analyzed data. CH set up the Cut &Tag analysis pipeline. OV, KU and LGM and AO interpreted the data. OV and AO wrote the manuscript. OV, CJ, OR, NC, LGM, KU and AO edited the manuscript.

## Data and materials availability

Cut&Tag data have been deposited in GEO under accession code GSE244768 (currently only accessible with the private token oxalksgoxdwbbyb). Source data are provided for cell tracking and Tbx6 profiles. All other data and raw microscopy data are available upon reasonable request. Python scripts are available on GitHub (https://github.com/EPFL-TOP/Tbx6_Oscillations).

