## Supplementary material for "Clock driven waves of Tbx6 expression prefigure somite boundaries"

**Supplementary Discussion 1:**

Here, we provide a description of the onset of the segmentation clock in zebrafish based on light-sheet timelapses of three embryos carrying the Her1-YFP transgenes starting around 30% epiboly. The Her1-YFP signal appeared about 20 minutes before the involution of cells at the blastoderm margin (data not shown), which marks the beginning of gastrulation at 50% epiboly (5.25 hpf), indicating that the onset of the segmentation clock occurs shortly before the start of gastrulation. The Her1-YFP signal quickly spread around the margin before travelling towards the animal pole in cells located in the epiblast (Fig. 1c-c'). A second wave of Her1-YFP signal traveled from the margin towards the animal pole when the embryo is at shield stage (6hpf). This wave took place all around the margin, both in the epi- and hypoblast (Fig. 1c-c'). Interestingly, cells in the hypoblast are advected towards the animal pole, in the same direction as the wave, while cells in the epiblast are advected in the opposite direction. This indicates that, at least this wave is kinematic and does not rely on cell advection. When the third wave started, the shield was now formed and devoid of Her1-YFP signal. Except for the shield, Her1-YFP was still expressed all around the margin but its expression was stronger on both sides of the shield (Fig. 1c-c'). This wave took place exclusively in the hypoblast. At around 75% epiboly, a fourth wave, seemingly restricted to the PSM, traveled a much longer distance than previous waves (Fig. 1c-c'). This is in agreement with results from in situ hybridization staining reporting a wave of *her1* expression travelling anteriorly at 70% epiboly[27]. As the fourth Her1-YFP wave arrived in the anterior PSM, a fifth wave started travelling from the posterior PSM (Fig. 1c').

Cell tracking revealed that cells forming the boundary B1 displayed on average 4 cycles of Her1-YFP expression as they were advected along the PSM (Fig. 1d). As each oscillation corresponds to a spatiotemporal wave of Her1-YFP expression, this suggests that the Mesp cell fate is instructed by the wave number 4, or less. As the first three waves of Her1-YFP expression only travel a short distance at the margin, more than two hours before the appearance of the first Mesp stripe, it is very likely that the fourth wave of Her1-YFP expression is the one instructing the Mesp cell fate of cells forming the first somite boundary.

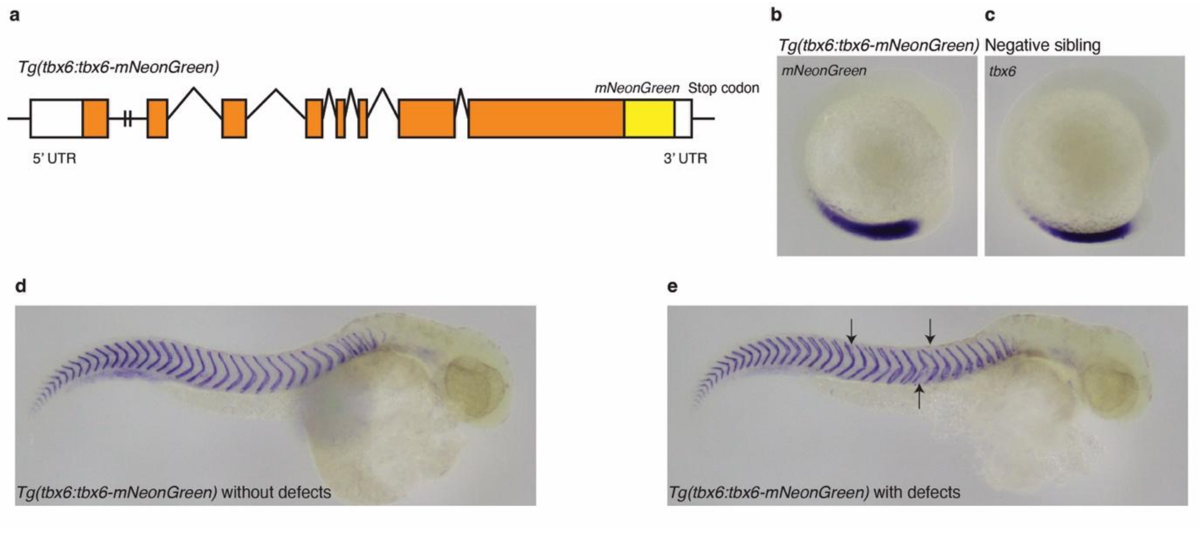


Extended Data Fig. 1: **Tg(*tbx6:tbx6-mNG*) recapitulates the endogeneous expression of *tbx6*. a,** the stop codon of the *tbx6* coding sequence was replaced with mNeonGreen. **b, c,** *In situ* hybridization staining for *mNeonGreen*, respectively *tbx6*, at 10-somite stage show similar patterns between Tg(*tbx6:tbx6-mNeonGreen)* and their negative siblings, respectively. **d, e,** *In situ* hybridization staining for *xirp2a* which marks the myotome boundaries show a normal segmentation phenotype (**d**, n = 12 embryos) or sporadic mild defects in (**e**, n = 14 embryos).

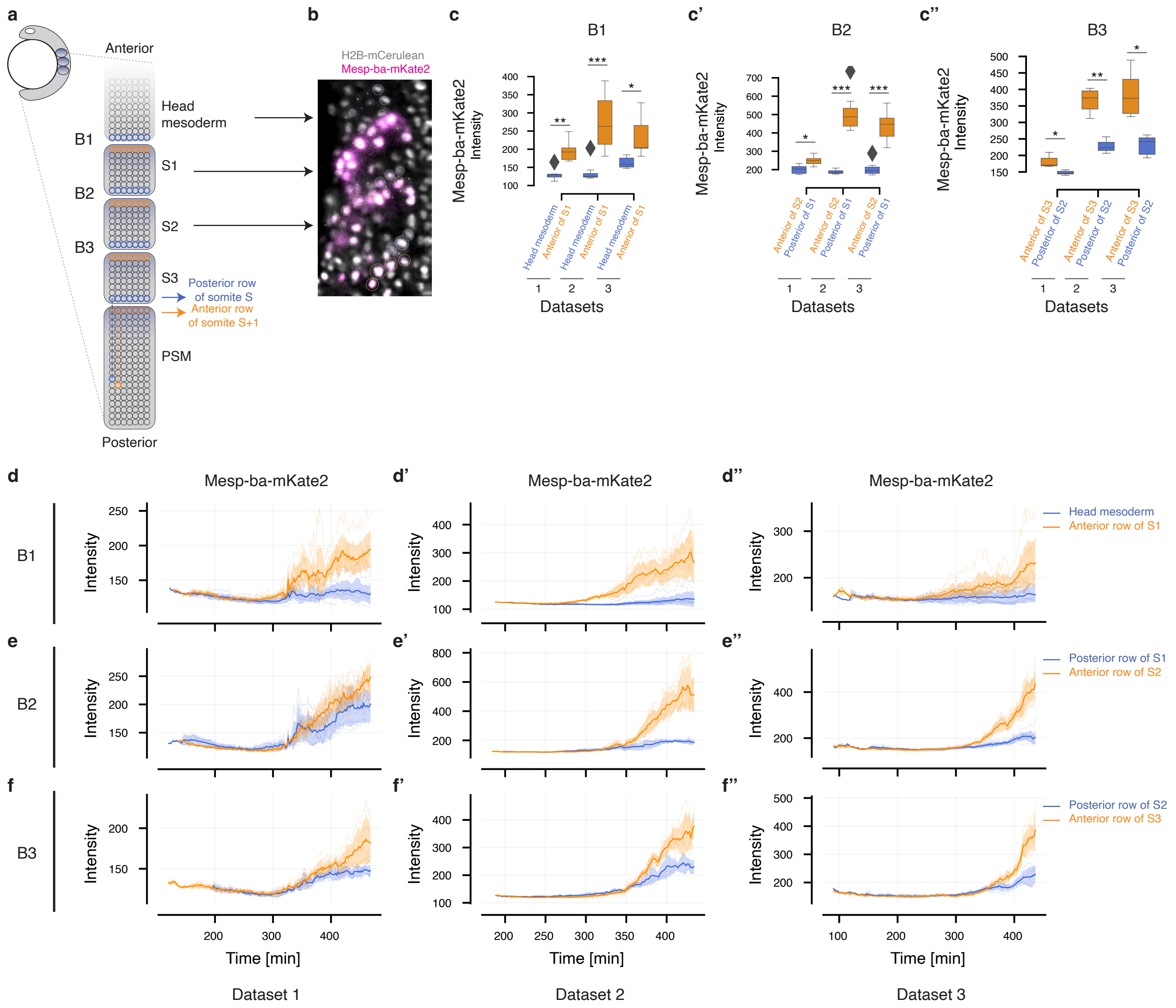


Extended Data Fig. 2: **Mesp-ba-mKate2 is expressed in the anterior part of somites. a,** Schematic of a zebrafish embryos during early somitogenesis. Somite boundaries form between Mesp-positive cells forming the anterior row of a somite (orange) and Mesp-negative cells forming the posterior row of the previous somite (blue). Cells forming somite boundaries were selected and back-tracked as they travelled along the PSM (blue and orange dashed lines). **b,** Snapshot from the Fiji plugin Mastodon of the first two somites of a zebrafish embryo. Boundary cells are identified using their H2BmCerulean and Mesp-ba-mKate2 signals and back-tracked in the PSM using their H2B-mCerulean signal. Circles represent the spots used for cell tracking. **c-c'',** Boxplots of Mesp-ba-mKate2 intensities of 3 Tg(*tbx6:tbx6-mNG;mesp-ba-mKate2)* embryos for cells forming the boundaries B1, B2 and B3 at the end of the traces shown in (**d-f''**)(n = 52, n = 41 and n = 31 cells from 3 embryos in **c**, **c'** and **c''**, respectively). Boundary cells located in the anterior row of a somite expressed significantly more Mesp-ba-mKate2 than boundary cells located in the posterior row. *P< 0.01, **P<0.001, ***P<0.0005. **d-f'',** Mesp-ba-mKate2 traces of the 3 Tg(*tbx6:tbx6-mNG;mesp-ba-mKate2)* embryos for cells forming the boundaries B1, B2 and B3.

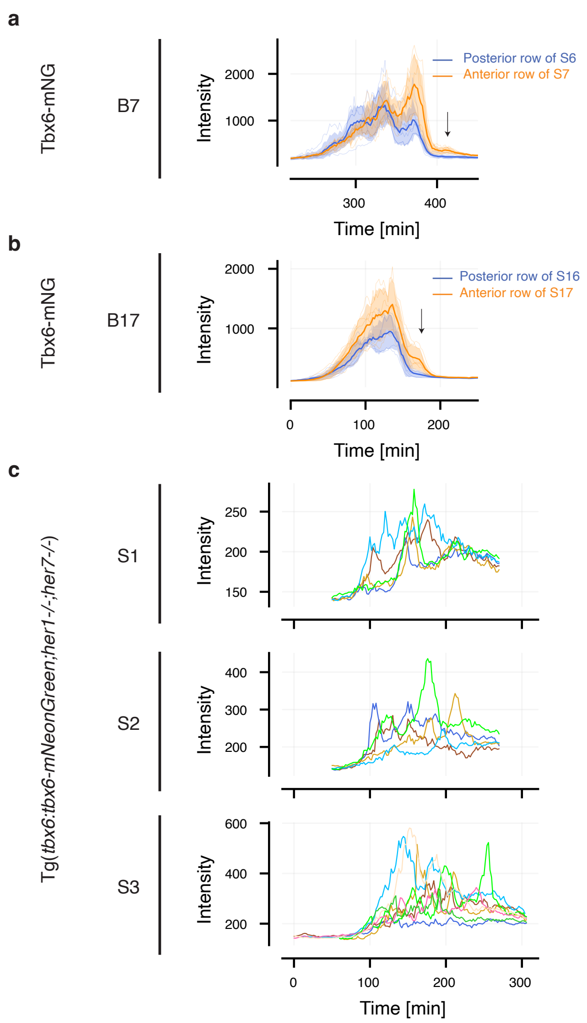


Extended Data Fig. 3: **Additional Tbx6-mNG traces. a, b,** Tbx6-mNG traces in cells forming the boundaries 7 and 17, respectively (n = 12 cells from 1 embryo and n=11 cells from another embryo, respectively). Traces from different embryos cannot be temporally aligned using the time of epiboly completion during mid-somitogenesis. Arrows indicate the last cycle of Tbx6 that distinguishes cells forming the anterior row of a somite (orange) and cells forming the posterior row of the previous somite (blue). **c,** Tbx6-mNG traces in cells located respectively in somites 1, 2 and 3 of a *her1;her7* double mutant (n =5, n = 5 and n = 8 cells from 1 embryo).

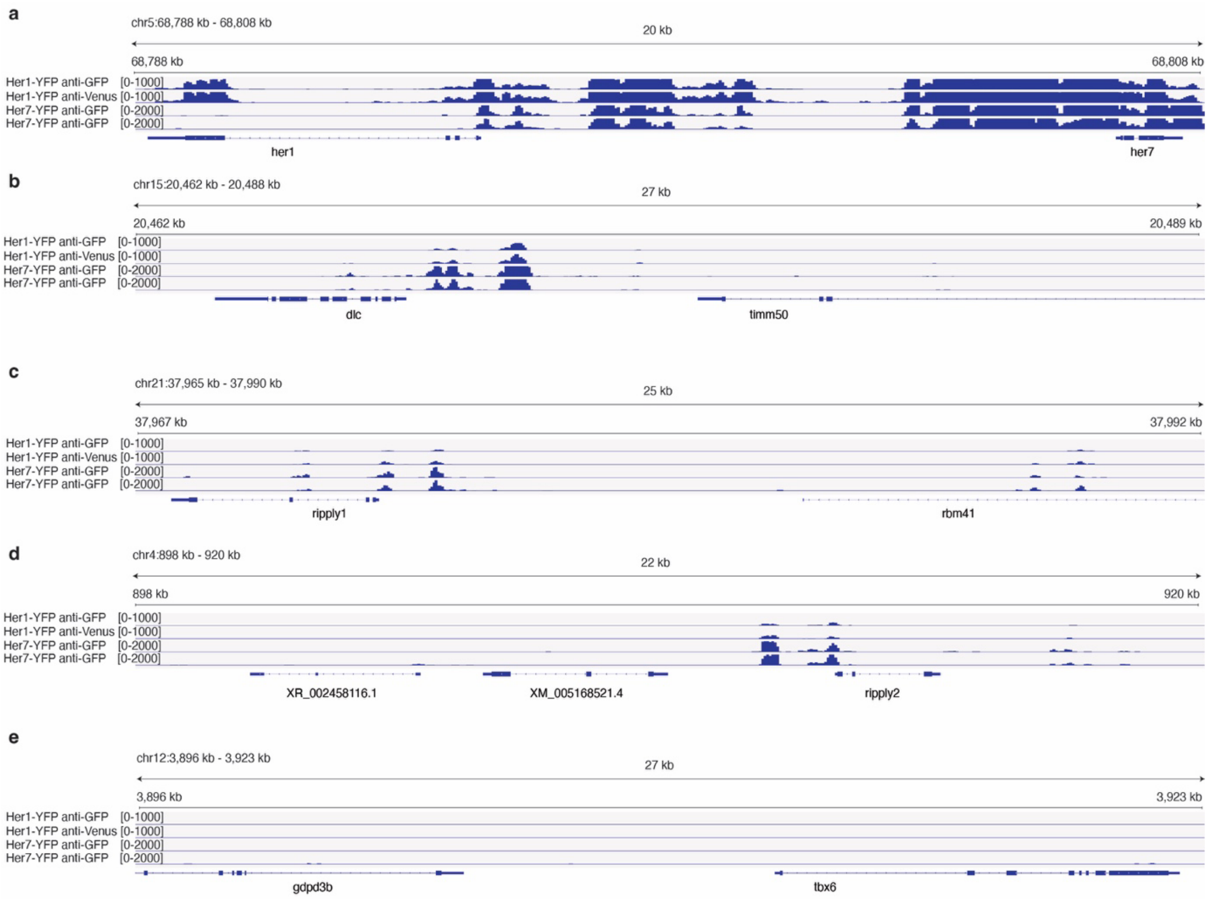


Extended Data Fig. 4: **Her1 and Her7 do not bind in the vicinity of the *tbx6* promoter. a-e,** Genomic binding landscapes provided by Cut&Tag of the fusion proteins Her1-YFP and Her7-YFP in the vicinity of *her1* and *her7* (**a**), *dlc* (**b**), *ripply1* (**c**), *ripply2* (**d**) and *tbx6* (**e**) coding regions. Each row corresponds to a biological replicate. Her1 and Her7 are bound to intergenic regions in the proximity of known and expected targets, such as their own chromosomal locus, *deltaC*[32,34], *ripply1*[25] and *ripply2*[25] (Extended Data Table 1 and 2), but no Her1 nor Her7 binding sites were found in the proximity of the *tbx6* gene.

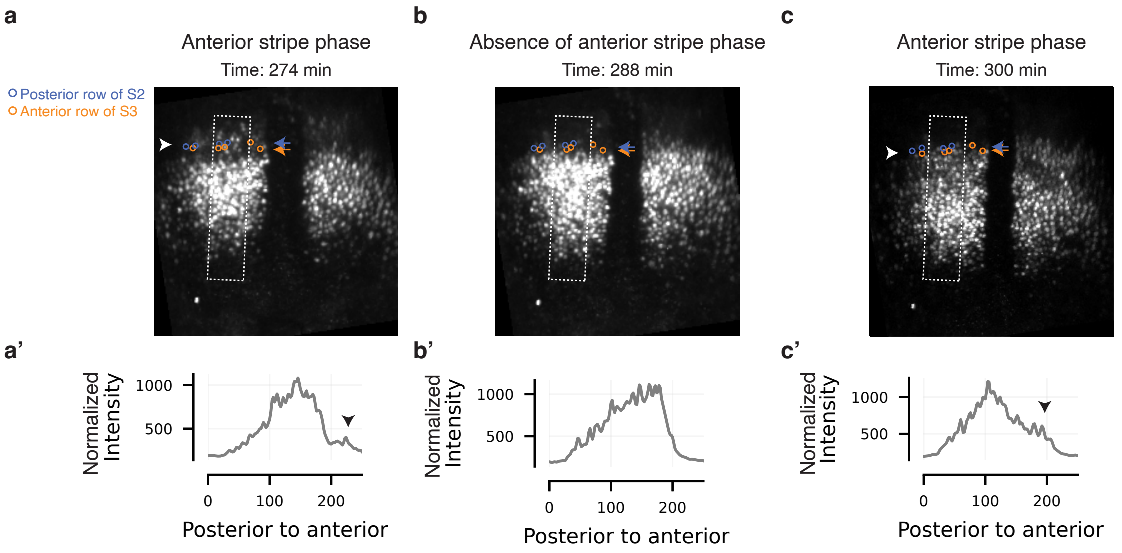


Extended Data Fig. 5**: Tbx6-mNG patterns and profiles match that of immunostainings against endogenous Tbx6**. **a, b, c,** Snapshots from a Tbx6-mNG timelapse at different timepoints (i. e. the "anterior stripe" phase, the next "absence of anterior stripe" and "anterior stripe" phases, respectively). Blue and orange circles mark the positions of cells located in the posterior row of somite 2 and cells located in the anterior row of somite 3, respectively. White boxes show the lines of interest used to plot the Tbx6 intensity profiles along the PSM (**a'-c'**). White arrowheads indicate the anterior stripe of Tbx6-mNG expression (**a**, **c**) and black arrowheads indicate the signature of this stripe on the intensity profiles (**a'**, **c'**)

| Chromosome | Peak start | Peak stop | Peak size | Proximal gene |
| --- | --- | --- | --- | --- |
| Ch5 | 68791966 | 68801274 | 9309 | *her1* |
| Ch5 | 68802800 | 68809533 | 6734 | *her7* |
| Ch15 | 20467327 | 20471678 | 4352 | *dlc* |
| Ch21 | 37972380 | 37972887 | 508 | *ripply1* |
| Ch4 | 910988 | 913327 | 2340 | *ripply2* |

**Extended Data Table 1: Cut&Tag of Her1-Venus.** Table listing a selection of relevant positions of significant peaks identified by Cut&Tag of Her1-Venus (see Methods).

| Chromosome | Peak start | Peak stop | Peak size | Proximal gene |
| --- | --- | --- | --- | --- |
| Ch5 | 68794962 | 68796302 | 1341 | *her1* |
| Ch5 | 68803185 | 68810009 | 6825 | *her7* |
| Ch15 | 20470627 | 20471515 | 889 | *dlc* |
| Ch21 | 37973727 | 37974213 | 487 | *ripply1* |
| Ch4 | 911024 | 912960 | 1937 | *ripply2* |

**Extended Data Table 2: Cut&Tag of Her7-Venus.** Table listing a selection of relevant positions of significant peaks identified by Cut&Tag of Her7-Venus (see Methods).

**Supplementary Video 1:** Maximum-intensity projection of a representative 3D timelapse of a zebrafish embryo carrying the Her1-YFP (yellow) and the Mesp-ba-mKate2 (magenta) transgenes and injected with H2B-mCerulean (cyan). Embryos were at the stage of 30% epiboly when the timelapse started. The time interval is 2 min.
**Supplementary Video 2:** Maximum-intensity projection of the Her1-YFP channel (16-color LUT) of an elliptically transformed version of Supplementary Video 1 (see Methods).
**Supplementary video 3:** Maximum-intensity projection of a representative 3D timelapse of a zebrafish embryo carrying the Tbx6-mNG (yellow) and the Mesp-ba-mKate2 (magenta) transgenes and injected with H2B-mCerulean (cyan). Embryos were at the stage of 30% epiboly when the timelapse started. The time interval is 2 min.
**Supplementary video 4:** Maximum-intensity projection of the Tbx6-mNG channel (16-color LUT) of an elliptically transformed version of Supplementary Video 3 (see Methods).
**Supplementary video 5:** Maximum-intensity projection of the Tbx6-mNG channel (16-color LUT) of a representative, elliptically transformed 3D timelapse of a *Tbx6-mNG*$^{+/-}$*;her1*$^{-/-}$*;her7*$^{-/-}$ zebrafish embryo. Tbx6-mNG is in yellow. The time interval is 2 min.
